# Altered cell wall hydroxycinnamate composition impacts leaf and canopy-level CO_2_-uptake and water-use in rice

**DOI:** 10.1101/2023.03.10.532111

**Authors:** Varsha S. Pathare, Rahele Panahabadi, Balasaheb V. Sonawane, Anthony Jude Apalla, Nouria Koteyeva, Laura E. Bartley, Asaph B. Cousins

**Author notes:** Author for correspondence: Varsha S. Pathare (;).

## Abstract

Cell wall properties can play a major role in determining photosynthetic carbon-uptake and water-use through impacts on mesophyll conductance (CO_2_ diffusion from substomatal cavities into photosynthetic mesophyll cells) and leaf hydraulic conductance (water movement from xylem, through leaf tissue to stomata). Consequently, modification of cell wall properties is proposed as a major path for improving photosynthesis and crop water-use efficiency. We tested this using two independent transgenic rice lines that overexpress the rice *OsAT10* gene (a “BAHD” CoA acyltransferase) which altered cell wall hydroxycinnamic acid content (greater *para*-coumaric acid and lower ferulic acid). Plants were grown under high and low water-levels and traits related to leaf anatomy, cell wall composition, gas exchange and hydraulics, plant biomass, and canopy-level water-use were measured. Alteration of hydroxycinnamic acid content led to significant decreases in mesophyll cell wall thickness (−14%), and increased mesophyll conductance (+120%) and photosynthesis (+22%). However, concomitant increases in stomatal conductance negated increased photosynthesis, resulting in no change in intrinsic water-use efficiency (ratio of photosynthesis/stomatal conductance). The leaf hydraulic conductance was also unchanged; however, the transgenics showed small, but significant increase in above-ground biomass (+12.5%), and canopy-level water-use efficiency (+8.8%; ratio of above-ground biomass/ water-used) and performed better under low water-level. Our results demonstrate that changes in cell wall composition, specifically hydroxycinnamic acid content, can increase mesophyll conductance and photosynthesis in C_3_ cereal crops like rice. However, attempts to improve photosynthetic water-use efficiency will need to enhance mesophyll conductance and photosynthesis whilst maintaining or decreasing stomatal conductance.

## Introduction

Water availability is a key factor limiting productivity and plant survival across both natural and managed ecosystems. Ongoing climate change will exacerbate these water limitations, further imposing constraints on crop production. Therefore, any future increases in crop production must be accomplished under existing or even lower water availability. This can be achieved by increasing plant water-use efficiency (WUE), defined as the amount of plant biomass produced per unit water used (Leakey et al., 2019). At the leaf-level, this means increasing WUE_i_ or the ratio of net photosynthetic CO_2_-uptake (A_net_) to stomatal conductance to water vapor (g_sw_). Attempts have been made to enhance WUE_i_ by lowering g_sw_. However, as A_net_ is positively associated with g_sw_, decreasing g_sw_ often decreases A_net_ (Leakey et al., 2019). Hence manipulation of A_net_ for improved WUE_i_ must happen independent of g_sw_. One possible means to achieve this is by enhancing mesophyll conductance to CO_2_ (g_m_) to supply a greater CO_2_ partial pressure (*p*CO_2_) at the site of Rubisco fixation for a given g_sw_ (Flexas et al., 2016; Leakey et al., 2019). In fact, g_m_ relates positively with A_net_ across diverse terrestrial plant groups (Flexas et al., 2012; Barbour and Kaiser, 2016; Pathare et al., 2020) and has been identified as the third most important factor limiting A_net_, besides stomatal and biochemical limitations (Gago et al., 2020). Also, because the CO_2_ diffusion pathways related to g_m_ are not the same as the pathways of water transpired out of stomata, an increase in g_m_ is expected to increase A_net_ without a concurrent increase in g_sw_. Consequently, modification of g_m_ through changes in leaf properties has been proposed as a key path for simultaneously improving crop A_net_ and WUE_i_ (Flexas et al., 2016; Leakey et al., 2019; Lundgren and Fleming, 2020). However, g_m_ in C_3_ species has been shown to coordinate with leaf hydraulic conductance (K_leaf_, the movement of water from xylem, through leaf tissue to stomata) in part due to the common pathway of CO_2_ and water movement inside leaves (Flexas et al., 2013; Xiong et al., 2017) but see (Loucos et al., 2017; Pathare et al., 2020). Studies also report a strong coupling of g_m_ and g_sw_ in C_3_ species, the mechanisms of which remain unclear (Hanba et al., 2004; Giuliani et al., 2013; Barbour and Kaiser, 2016). This suggests that increases in g_m_ may lead to simultaneous increases in K_leaf_ and g_sw_ thus hampering our efforts to improve WUE_i_ through modification of g_m_. Consequently, identification of leaf traits that enhance g_m_, whilst maintaining K_leaf_ and g_sw_, will be a critical step towards exploiting g_m_ for improving crop A_net_ and WUE_i_.

Both g_m_ and K_leaf_ are complex traits influenced by several leaf structural, anatomical, and biochemical properties (Evans et al., 2009). Among these, mesophyll surface area exposed to intercellular air spaces (S_mes_), chloroplast coverage of S_mes_ (S_c_) and mesophyll cell wall (CW) properties (Xiong and Flexas, 2018; Pathare et al., 2020; Evans, 2021; Flexas et al., 2021) have been identified as the strongest determinants of g_m_ across diverse plant groups. Greater S_mes_ is expected to increase g_m_ by increasing the surface area for diffusion of CO_2_ inside photosynthetic mesophyll cells; whereas, greater S_c_ is expected to increase g_m_ by decreasing the path length for CO_2_ diffusion inside the chloroplast (Evans et al., 2009). S_mes_ is also expected to play a role in mediating the coordination of g_m_ and K_leaf_, primarily the outside xylem conductance (K_ox_), by increasing the surface area for evaporation of water (Xiong et al., 2017). Mesophyll CWs are another key leaf anatomical feature that exert a major influence on g_m_ and hence A_net_ that is equal to or even greater in magnitude than S_mes_ (Evans, 2021; Flexas et al., 2021).

CWs are complex and dynamic biological structures that affect several critical plant functions (Keegstra, 2010), including the exchange of and CO_2_ and water inside leaves (Evans, 2021; Flexas et al., 2021). The structure and composition of CWs vary greatly depending on the plant group (e.g., dicot versus monocot), cell types and developmental stage (Sarkar et al., 2009). Photosynthetic mesophyll CWs are mainly composed of variable proportions of cellulose, several matric polysaccharides, structural proteins and other minor components (Sarkar et al., 2009; Cosgrove and Jarvis, 2012). The structure, organization and interactions of these components results in a web of nanometric and micrometric pores that are filled with apoplastic liquid. These pores allow the movement of CO_2_ and water between the intercellular air spaces and mesophyll cells (Evans et al., 2009). According to some estimates, mesophyll CWs can account for > 50% of the total resistance to CO_2_ diffusion from the substomatal cavities into the mesophyll cells. This resistance (r_w_) can be expressed as 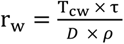 (Evans et al., 2009; Evans, 2021), where, T_CW_ is the CW thickness, τ is the tortuosity of the liquid path through the walls, *D* is the diffusivity of CO_2_ in apoplastic liquid and *ρ* is the porosity of the wall. Previous studies demonstrate that thicker T_CW_ increases the apparent diffusion pathlength, thus decreasing g_m_ (Evans, 2021; Flexas et al., 2021). In addition, a few recent studies based on diverse trees and herbaceous dicot species have proposed a role for CW components in determining g_m_ by affecting T_CW_, porosity and tortuosity (Weraduwage et al., 2016; Roig-Oliver et al., 2020; Flexas et al., 2021; Roig-Oliver et al., 2022). For instance, greater cellulose and hemicellulose content are expected to reduce g_m_ by increasing tortuosity. Whereas a relatively higher pectin content and a greater pectin to cellulose and hemicellulose ratio is expected to increase CW hydrophilicity and effective porosity to CO_2_, thus increasing g_m_ (Roig-Oliver et al., 2020; Flexas et al., 2021). Furthermore, lignin deposition is expected to decrease CW porosity and g_m_ (Roig-Oliver et al., 2020). In addition to the above major components, CW-bound phenolic compounds are also expected to affect g_m_ and A_net_ by influencing the chemical tortuosity or any chemical interactions of CO_2_ with major CW components (Flexas et al., 2021). The above studies provide significant information about how and why different CW components can affect CO_2_ diffusion inside mesophyll cells and highlight their importance for improving crop g_m_ and A_net_ (Evans, 2021; Flexas et al., 2021). However, most studies have focused on trees and herbaceous dicots, whereas impacts of different CW components on g_m_ and A_net_ have rarely been studied on grasses, which represent many of the cereal crops (Hura et al., 2012; Ellsworth et al., 2018).

Grass CWs differ significantly from dicots in structure and composition (Vogel, 2008). The primary CWs of dicots, which we expect to be present in stomatal-facing mesophyll cells, mainly consist of a cellulose-xyloglucan network embedded in a pectin matrix. In contrast, grass primary walls are low in xyloglucan and pectin. Instead, xylans (glucuronoarabinoxylan) and mixed-linkage glucan are the principal polymers that interact with cellulose microfibrils in grass primary CW(Cosgrove and Jarvis, 2012). Another feature that distinguishes grass CW from those of dicots is the incorporation of hydroxycinnamates like ferulic acid (FA) and *para*-coumaric acid (*p*-CA) onto arabinoxylan (Chandrakanth et al., 2023). *p*-CA is also esterified to lignin present in secondary CW thickenings of leaf vascular bundles, and, at lower levels, FA has also been found on lignin of various species (Ralph, 2010; Karlen et al., 2016). These hydroxycinnamate moieties affect CW properties, including the aggregate property of cellulose enzymatic digestibility, which may relate to wall porosity or “looseness” in grasses (Bartley et al., 2013; Tian et al., 2021). A recent study on rice showed that mixed-linkage glucan, a polysaccharide unique to grasses, plays key role in determining CW porosity, g_m_ and A_net_ (Ellsworth et al., 2018). However, the impacts of most grass CW components on g_m_ and A_net_ remain unknown. A greater understanding of how different grass CW components affect the movement of CO_2_ inside photosynthetic mesophyll cells will be critical for making targeted changes in CW properties to improve g_m_ and A_net_ of cereal crops.

Mesophyll CW properties are also expected to impact K_leaf_ or the water movement from the xylem through leaf tissue to stomata (Sack and Holbrook, 2006; Buckley et al., 2015). K_leaf_ can be partitioned into two major components: inside and outside xylem pathways (K_x_ and K_ox,_ respectively). While K_x_ and traits (like vein density) influencing it are well characterized, K_ox_ remains poorly understood. The outside-xylem pathways of water movement are highly complex as water travels via apoplastic (CWs) and/or symplastic (cell-to-cell via plasmodesmata) routes and involves multiple leaf tissues including bundle sheath, mesophyll, and epidermis (Buckley et al., 2015). While the relative importance of apoplastic versus symplastic routes in determining K_ox_ remains controversial, studies suggest that the apoplastic route via CWs is a major contributor to water movement, influenced by CW properties like T_CW_. For instance, using model simulations, Buckley *et al*. (2015) demonstrated that an increase in T_CW_ can increase the outside-xylem apoplast path length, thereby increasing K_ox_ and hence K_leaf_. In addition to T_CW_, CW composition is also expected to influence K_leaf_ (Sack and Holbrook, 2006). However, despite the importance of K_leaf_ in determining A_net_ and g_sw_, little is known about how leaf traits like CW properties influence K_leaf_.

Toward understanding the influence of CW properties on g_m_ and K_leaf_ in a grass, we used two independent transgenic rice lines overexpressing *OsAT10* gene (a BAHD Acyltransferase) to test the impacts of CW modification on traits related to cellular, leaf and plant-level CO_2_-uptake and water-use, including g_m_, A_net_ and K_leaf_. Overexpression of the *OsAT10* gene alters rice CW hydroxycinnamic acid content resulting in greater *p-*CA and lower FA content and thus increasing degradability of rice biomass (Bartley et al., 2013). Previous studies on triticale genotypes also suggest the potential role of these hydroxycinnamic acids in determining productivity and drought tolerance (Hura et al., 2012). Here, we assessed whether modification of CW hydroxycinnamic acids content affects g_m_ and K_leaf_ and whether these cellular and leaf-level changes scale to influence plant-level biomass and WUE. We demonstrated that altered CW hydroxycinnamate composition increased g_m_ and A_net_, but not K_leaf_. Still, the transgenic plants observed an overall increase in tolerance to water restriction.

## Methods

### Plant material and growth conditions

Previously characterized *Oryza sativa* spp. *japonica* and transgenics (*ZmUbi_pro_-AT10* and *OsAT10-D1*) overexpressing the gene *OsAT10 (LOC_Os06g39390)* were used in the current study (Bartley et al., 2013).The *ZmUbi_pro_-AT10* line (Line 7-1), in which the rice *AT10* coding sequence is driven by the *Zea mays Ubiquitin1* promoter, and a negative segregant (Line 7-5) are in the Kitaake cultivar and the activation-tagged construct (*OsAT10-D1*, Line 21) and a negative segregant (Line 17) are in the Dongjin cultivar. Overexpression of *OsAT10* in rice was previously reported to increase FA and decrease *p*-CA, and increase digestibility of rice, without noticeable impacts on the health and vegetative development of transgenic plants. Also, no significant changes in lignin content and composition were reported, though an increase in CW glucose content was noted (Bartley et al., 2013).

Seeds for WT and transgenics were germinated on wet filter paper in a Petri plate for four days and then transplanted to trays containing Sunshine Mix LC-1 soil (Sun Gro Horticulture, Agawam, MA, USA) mixed with turface (ratio of 3: 1 in volume). Ten days after germination, seedlings were transplanted into 3-L free-drainage pots in climate-controlled growth chambers (Biochambers, #GRC-36, Winnipeg, MB, Canada). The photoperiod was 14 h, including two h ramps at the beginning and end of the light period. Temperatures during light and dark phase were maintained at 26 and 22°C, respectively, and relative humidity levels were at 70%. The light was emitted by 400W metal halide and high-pressure sodium lamps with a maximum photosynthetic photon flux density (PPFD) of c. 1000 µmol photons m^−2^ s^−1^.

This experiment consisted of two water levels, high-water (HW) or 90% of pot capacity and low-water (LW) or 50% of pot capacity and 4 lines (2 WT and 2 transgenics) with 6 replicates for each treatment. Before transplanting, all the pots were weighed, then filled with potting mix and weighed again to get empty pot and pot plus potting mix weights. These pots were saturated with water and allowed to drain overnight and weighed the next morning to get the 100% pot capacity weight. Ten-day old seedlings were then transplanted to pots (one seedling in each pot). Water treatments were initiated by withholding irrigation until the pots reached the desired 90% and 50% pot capacity weights for HW and LW treatments respectively (which were achieved in 4 days for HW pots and 11 days for LW pots). For the remainder of the experiment pots were maintained at HW and LW conditions by weighing each day and watering to the target weights. Pots and their lateral holes were covered with plastic to minimize evaporation. Daily transpiration was measured as the daily difference in pot weight minus soil evaporation (measured as the difference in weight of a covered pot without a plant in each treatment group). Total water used (g) was calculated at the end of the experiment by summing the daily transpiration values. Pots were randomly moved around each day when pots were weighed and watered.

### Gas exchange and mesophyll conductance measurements

Simultaneous measurements of net photosynthetic rates (A_net_; µmol m^−2^ s^−1^), stomatal conductance to water vapor (g_sw_; mol water m^−2^ s^−1^), intercellular *p*CO_2_ (C_i_, Pa) and photosynthetic C-isotope discrimination was conducted using a Li-6800 portable photosynthesis system (LI-6800, Li-Cor, Lincoln, USA) coupled to a tunable diode laser absorption spectroscope (TDLAS, model TGA 200A; Campbell Scientific, Logan, UT, USA). Measurements were conducted 45-52 days after germination on the top-most fully expanded leaves around mid-day (9:30-14:00) for each line and water treatment. Multiple non-overlapping leaves were placed across the Li-6800 chamber and were allowed to adjust for at least 30 min or until stable values of A_net_ and g_sw_ were achieved. The CO_2_ sample was maintained at *p*CO_2_ of ~ 37 Pa, leaf temperature at 26°C, and relative humidity at 55-60%. Data for isotopologs of CO_2_ and H_2_O and physiological parameters (A_net_, g_sw_, C_i_) were collected and averaged over the next 20–30 min for c. 8–12 cycles of TDLAS with the Li-6800 was set to log data only when the TDLAS analyzed the Li-6800 sample line. 5-6 biological replicates per treatment were measured. Leaf-level intrinsic water-use efficiency (WUE_i_, μmol CO_2_ mol_−1_ water) was calculated as A_net_/g_sw_. Mesophyll conductance to CO_2_ (g_m_, µmol CO_2_ m^−2^ s^−1^ Pa^−1^) was estimated using the _13_C method for C_3_ species as detailed in previous studies (Ellsworth et al., 2018; Sonawane and Cousins, 2019).

After measurement of physiological parameters and isotopologs, photosynthetic CO_2_ response curves (A_net_-C_i_ curves) were measured on the same leaves to determine photosynthetic capacities. A_net_-C_i_ response curves were measured with thirteen different steps of *p*CO_2_ (5, 10, 14, 19, 23, 28, 37, 51, 65, 93, 111, 139, 167 Pa) while maintaining saturating light conditions (photon flux density of 1500 μmol photons m^−2^ s^−1^), 55 – 65% relative humidity and leaf temperatures of 26°C and allowing a stabilization time of 2 - 3 minutes after each step change in *p*CO_2_. During the A_net_-C_i_ measurements, *p*CO_2_ in the cuvette was controlled in the Li-COR sample line. 5-6 replicate plants per treatment condition were used to measure A_net_-C_i_ curves. A_net_-C_i_ curves were then fitted using the biochemical model of photosynthesis for C_3_ species (Farquhar et al., 1980), to obtain kinetic coefficients associated with rates of maximum carboxylation (V_cmax_; μmol CO_2_ m_−2_ s_−1_) and electron transport (J_max_; μmol electron m_−2_ s_−1_). While deriving V_cmax_ and J_max_, we used g_m_ values estimated in the current study for each line and treatment condition. Maximum photosynthetic rates (A_max_, μmol CO_2_ m_−2_ s_−1_) were measured at saturating light of ~ 1,200 μmol photons m_−2_ s_−1_ and *p*CO_2_ of 1500 μmol CO_2_ mol_−1_ air. After gas exchange measurements, leaves were harvested immediately and processed to analyze CW composition, leaf N content, and leaf anatomy.

### Estimation of stomatal, mesophyll and biochemical limitations to photosynthesis

We estimated limitations to light-saturated CO_2_ assimilation rates that primarily occur through stomatal restrictions to the diffusion of CO_2_ into intracellular leaf spaces (L_s_), mesophyll restrictions to the diffusion of CO_2_ into the site of fixation by Rubisco (L_m_), and due to the biochemistry of CO_2_ fixation at the chloroplast (L_b_) as described previously (Jones, 1985; Grassi and Magnani, 2005). In this approach, the total photosynthetic limitation is divided into the relative limitations of stomata (L_s_), mesophyll (L_m_), and biochemistry (L_b_):

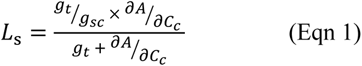

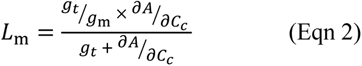

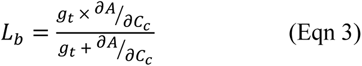

where *g_t_* is the total conductance to CO_2_ from the leaf surface to the sites of fixation at Rubisco:

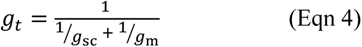

and, ∂*A*/∂*C_c_* is the partial derivative of net CO_2_ assimilation (A_net_) for the relative change in CO_2_ concentration at the Rubisco (C_c_) derived from the initial slope of A_net_ - C_c_ curve as described earlier (Pathare et al., 2017). For this, we used *plantecophys* R-package (Duursma, 2015) to estimate C_i_ at the transition and compensation point of A_net_-C_i_ curve, and respective C_c_ values were calculated using measured g_m._

### Analysis of leaf CW composition, N content and anatomical traits

For CW composition and N content analysis, newly expanded, fully emerged leaves used for gas exchange measurements were harvested for WT and transgenic lines exposed to HW and LW treatments. Leaf samples were dried in a hot air oven at 55°C for 72 hours. Analysis of CW components, that is, ferulic acid (FA; µg/mg), para-coumaric acid (*p*-CA; µg/mg) and lignin (%) was performed as indicated previously for rice (Bartley et al., 2013; Karlen et al., 2016). Briefly, leaves were milled, and de-starched alcohol insoluble residues were prepared. This CW prep was treated first with weak acid (50 mM trifluoroacetate) to release matrix polysaccharide-associated phenolates and then both the CW pellet and acid supernatant were treated with base to cleave the esterified FA and *p-*CA, which were quantified by high-performance liquid chromatography. Leaf nitrogen (N) content was measured using a Eurovector elemental analyzer and expressed on a leaf area basis (N_area_; g m^−2^). Leaf mass per area (LMA; mg cm^−2^) was also determined using the gas exchange leaves as a ratio of dry leaf mass to leaf area.

Leaf anatomical traits associated with g_m_ and K_leaf_ were measured on WT and transgenic lines only for HW treatment on topmost fully expanded leaves (45-52 days after germination). Light and transmission and scanning electron microscopy techniques were used to measure leaf thickness (µm), length of mesophyll CWs exposed to intercellular air spaces (IAS; µm), mesophyll CW thickness (T_CW_; µm) and bundle sheath CW thickness (BS_CW_; µm). The details of sample preparation for light and electron microscopies, measurements and calculations are presented in our previous study (Pathare et al., 2020). Briefly, light microscopy images of leaf cross sections were used to measure the length of mesophyll CWs exposed to intercellular air spaces (IAS) using 10-15 different fields of view for each leaf (*n* = 4) taken at x 50 and x 100 magnifications. The mesophyll surface area exposed to intercellular air spaces (S_mes_; µm_2_ µm^−2^) was calculated from measurements of the total length of mesophyll CWs exposed to IAS and width of section analyzed using the equation from Evans et al., (1994) with curvature correction factor (F) of 1.34. Whereas the length of chloroplasts exposed to the IAS (S_c_; µm_2_ µm^−2^) was calculated using the TEM images taken randomly at 10-15 fields of view for each leaf (*n* = 4). T_CW_ and BS_CW_ were measured from TEM micrographs using at least 15 images for each leaf (*n* = 4). Light microscopy images of leaf cross sections were used to determine leaf thickness (µm) as the average values measured over the vascular bundles and the bulliform cells (*n* = 4). For interveinal distance (IVD; µm), the distance from the center of one vein to the other was measured.

### Measurement of Leaf hydraulic parameters

Measurements to obtain leaf water potential (Ψ_leaf_; MPa), leaf hydraulic conductance (K_leaf_; mmol m^−2^ s^−1^ MPa^−1^), leaf hydraulic conductance inside xylem (K_x_; mmol m^−2^ s^−1^ MPa^−1^) and outside xylem conductance to water (K_ox_; mmol m^−2^ s^−1^ MPa^−1^) were made on the youngest fully expanded leaves from an individual plant as described previously (Sonawane et al., 2021) only for the *Ubi_pro_-AT10* and corresponding negative segregant lines. The K_leaf_ and K_x_ measurements were performed at leaf temperatures of 26°C. During K_leaf_ measurements, the incident light on the leaf was 1200 μmol photons m^−2^ s^−1^. About five or six plants for each *Ubi_pro_-AT10* line and treatment combination were used (*n* = 5-6).

### Measurement of biomass and WUE_canopy_

Sixty days after germination, all plants were harvested and separated into the stem (non-transpiring tissue) and leaves (transpiring tissue) and dried in a ventilated drying oven at 55°C for a week before weighing to measure the above-ground biomass (AGB = stem biomass + leaf biomass; g) (*n* = 5-6). Leaf width (cm) was measured at the time of harvest on 10 different leaves per plant (*n* = 5-6). AGB and total water used during the duration of the experiment were used to calculate canopy-level water-use efficiency (WUE_canopy_ = g dry AGB/g water used).

## Statistical analyses

Statistical analyses and data visualization were performed using R software (v 4.1.0, R Foundation for Statistical Computing, Vienna, Austria). Normality and equal variances were tested, and when necessary square root or log transformations were used to improve the data homoscedasticity. Two-way ANOVA was performed for all the measured traits (except leaf anatomical traits) with line and water-level as the main effects using the *aov* function in R (R Core Team, 2018). Results of two-way ANOVA are given in Tables 1-3 for *Ubi_pro_-AT10* lines and Table S1 for *OsAt10-D1* lines. One-way ANOVA with *post-hoc* Tukey’s test was used to examine differences in measured traits among the four line and water treatment combinations, that is, WT and transgenic subjected to HW treatment and WT and transgenic subjected to LW treatments. Results for *post-hoc* Tukey’s test are given in Fig. 2–7 for *Ubi_pro_-AT10* lines and Fig. S1-S3 for *OSAT10-D1* lines. For the one-way ANOVA, *P*-values ≤ 0.05 were considered statistically significant and *P*-values ≤ 0.1 as marginally significant.

**Table 1:** Summary of two-way ANOVA P-values and F-values for the leaf CW composition, physiology and biochemical traits associated with CO_2_-uptake and water-use measured on the rice *Ubi_pro_-AT10* WT and transgenic lines at 50-60 days’ time point. *P*-values ≤ 0.1 are highlighted in bold. *P*-values ≤ 0.05 were considered as statistically significant and *P*-values ≤ 0.1 were considered as marginally significant.

| Abbreviations | Traits | Genotype |  | Water |  | Genotype x Water |  |
| --- | --- | --- | --- | --- | --- | --- | --- |
|  |  | *P*-value | *F*-value | *P*-value | *F*-value | *P*-value | *F*-value |
| FA-Xylan | Ferulic acid on Xylan | < **0.001** | 117.832 | **0.043** | 4.849 | **0.004** | 11.288 |
| FA-Lignin | Ferulic acid on Lignin | < **0.001** | 26.494 | **0.048** | 4.562 | 0.142 | 2.380 |
| *p*-CA-Xylan | Para-coumaric acid on Xylan | < **0.001** | 204.456 | 0.193 | 1.849 | 0.118 | 2.735 |
| *p*-CA-Lignin | Para-coumaric acid on Lignin | < **0.001** | 24.647 | **0.008** | 9.022 | 0.614 | 0.265 |
| Lignin | Lignin | 0.251 | 1.422 | 0.401 | 0.743 | 0.318 | 1.061 |
| A<sub>net</sub> | Net CO<sub>2</sub> assimilation rates | < **0.001** | 37.20 | 0.102 | 2.94 | 0.605 | 0.28 |
| Δ<sup>13</sup>C-g<sub>m</sub> | Mesophyll conductance to CO<sub>2</sub> | **0.001** | 16.66 | **0.003** | 11.44 | 0.315 | 1.06 |
| g<sub>sw</sub> | Stomatal conductance to water | **0.003** | 11.085 | **0.035** | 5.096 | 0.334 | 0.981 |
| Δ<sup>13</sup>C-g<sub>m</sub>/g<sub>sw</sub> | Ratio of mesophyll to stomatal conductance | **0.006** | 9.656 | **0.013** | 7.400 | 0.222 | 1.590 |
| C<sub>i</sub>/C<sub>a</sub> | Ratio of intercellular to atmospheric CO<sub>2</sub> partial pressure | 0.365 | 0.861 | 0.150 | 2.243 | 0.657 | 0.204 |
| WUE<sub>i</sub> | Intrinsic water-use efficiency (A<sub>net</sub>/g<sub>sw</sub>) | 0.303 | 1.12 | 0.111 | 2.78 | 0.619 | 0.26 |
| V<sub>cmax</sub> | Rates of maximum carboxylation | 0.957 | 0.00 | 0.525 | 0.42 | 0.271 | 1.29 |
| J<sub>max</sub> | Rates of electron transport | **0.003** | 11.57 | **0.090** | 3.19 | 0.445 | 0.61 |
| A<sub>max</sub> | Maximum photosynthetic rates | < **0.001** | 22.14 | **0.017** | 6.80 | 0.560 | 0.35 |
| L<sub>s</sub> | Stomatal limitations to A<sub>net</sub> | **0.015** | 7.10 | 0.116 | 2.71 | 0.420 | 0.68 |
| $L_m$ | Mesophyll limitaions to $A_{net}$ | **< 0.001** | 507.30 | **< 0.001** | 309.95 | **0.003** | 11.22 |
| $L_b$ | Biochemical limitaions to $A_{net}$ | **< 0.001** | 175.17 | **< 0.001** | 63.96 | **< 0.001** | 33.31 |
| $K_{leaf}$ | Leaf hydraulic conductance | 0.20 | 1.76 | **< 0.001** | 11.31 | 0.18 | 1.95 |
| $K_{ox}$ | Outside-xylem conductance to water | **0.09** | 3.27 | **0.01** | 7.83 | 0.60 | 0.29 |
| $K_x$ | Xylem conductance to water | 0.12 | 2.62 | **0.10** | 3.08 | 0.23 | 1.53 |
| Leaf water potential | Leaf water potential | 0.49 | 0.50 | 0.97 | 0.00 | 0.15 | 2.22 |

## Results

To ensure that any observed effects were not due to other genetic modifications independent of *AT10*-over expression, we used two distinct rice CW transgenics in this study (*ZmUbi_pro_-AT10* (hereafter, *Ubi_pro_-AT10*, and *OsAT10-D1*). Overall, we observed similar compositional, physiological, and anatomical changes in *Ubi_pro_-AT10* transgenics compared to the activation tagged *OsAT10-D1 (OsAT10-Dominant 1)*. Grass *Ubi* promoters generally have strong foliar expression, whereas the native *AT10* gene has low expression in leaves compared to other related BAHDs (Chandrakanth et al., 2023). Thus, we focused on the *Ubi_pro_-AT10* lines and measured a larger battery of traits, whereas, for *OsAT10-D1* only important physiological and structural traits were measured. Data for *Ubi_pro_-AT10* lines is included in the main document (Fig. 1 to Fig. 7). For *OsAT10-D1* transgenics, leaf anatomy data has been presented as the main figure (Fig. 1), while other data is given in the supplement (Fig. S1 to S3 and Table S1). Generally, physiological effects in the *Ubi_pro_-AT10* lines were more pronounced than for the *OsAT10-D1* lines.

**Figure 1.**
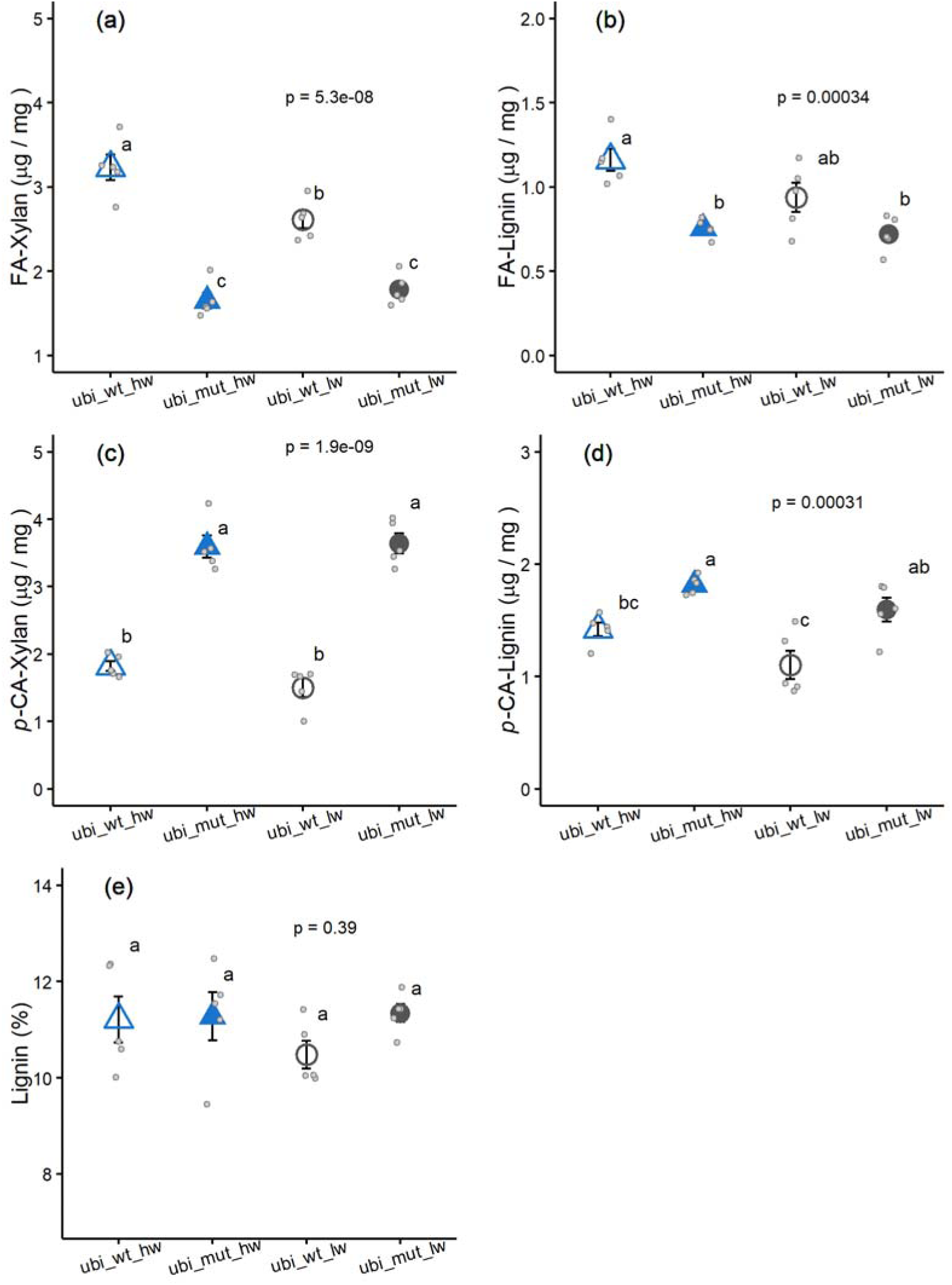
Leaf cell wall composition for *Ubi_pro_-AT10* wild type (ubi_wt, open symbols) and transgenic (filled symbols) measured under high water (hw, blue triangles) and low water (lw, gray circles). (a) Ferulic acid content on xylan (FA-Xylan); (b) Ferulic acid content on lignin (FA-Lignin); (c) *para*-coumaric acid content on xylan (*p-*CA-Xylan); (d) *para*-coumaric acid content on lignin (*p*-CA - Lignin); (e) lignin content. *P*-values from one-way ANOVA and post-hoc Tukey’s test letters are shown. Values indicate mean ± SE (*n*=5-6) along with replicate points (small open circles).

### Changes in CW composition

Since the CW composition of leaves of the developmental stage and growth conditions examined had not been reported previously for these transgenic lines, we evaluated the differences in leaf CW composition between the WT and transgenics growing under HW and LW treatments (Fig. 1, Table 1). There was a significant genotype effect on FA and *p*-CA content (*P* < 0.001, Table 1, Fig. 1a-d) for the *Ubi_pro_-AT10 lines*. The *Ubi_pro_-AT10* transgenics showed significantly lower FA-xylan and FA-lignin (~40 and 30% resp. Fig. 1a, b) but higher *p*-CA-xylan and *p*-CA-lignin (~100 and 150% resp. Fig. 1a, b) compared to WT. We also observed a significant genotype x water effect (*P* = 0.004, Table 1) for FA-xylan, wherein, *Ubi_pro_-AT10* transgenics showed greater decrease in FA-xylan content under HW treatment (44%) compared to LW treatment (30%). Whereas *Ubi_pro_-AT10* transgenics did not differ significantly from WT in terms of lignin content (*P* > 0.1, Table 1). Similar differences were observed between *OsAT10-D1* transgenics and WT (Fig. S1, Table S1). The *OsAT10-D1* transgenics showed significantly lower FA-xylan and FA-lignin (~53 and 59% resp. Fig. S1a, b) but higher *p*-CA-xylan and *p*-CA-lignin (~180 and 80% resp. Fig. S1a, b) compared to WT.

### Changes in leaf anatomical traits

We assessed changes in key leaf anatomical traits in the WT and transgenic plants growing under HW treatment. Both *Ubi_pro_-AT10* and *OsAT10-D1* transgenics exhibited lower mesophyll CW thickness (~13-15%) compared to WT plants, with the differences in CW thickness between WT and transgenics being statistically significant for *Ubi_pro_-AT10* (*P* < 0.05, Fig. 2a) and marginally significant for *OsAT10-D1* genotypes (*P* < 0.1, Fig. 2a). Whereas we did not observe statistically significant differences in BS_CW_ between WT and transgenics in both *Ubi_pro_-AT10* and *OsAT10-D1* lines (Fig. 2b). Furthermore, we observed a marginally significant decrease in S_mes_ (12%) in *Ubi_pro_-AT10* transgenics compared to WT (*P* = 0.062), but not in the *OsAT10-D1* transgenics (Fig. 2c). Other leaf anatomical traits, like S_c_, leaf thickness and IVD did not differ significantly between the WT and transgenic plants for both *Ubi_pro_-AT10* and *OsAT10-D1* lines (*P* > 0.1, Fig. 2d-f).

**Figure 2.**
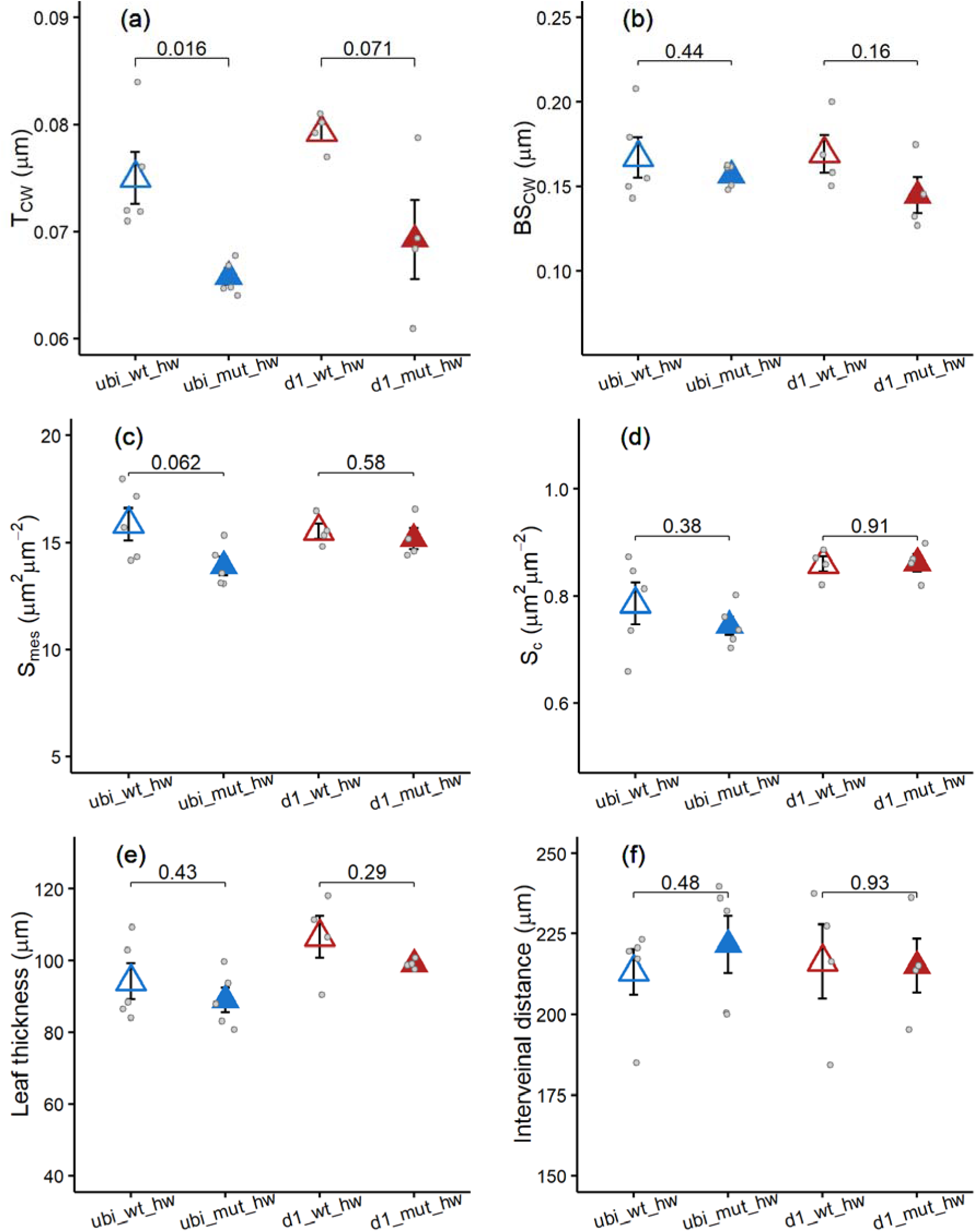
Leaf anatomical traits for rice genotypes: *Ubi_pro_-AT10* wild type (ubi wt; blue open triangles) and transgenic (ubi_mut; blue filled triangles) and *OsAt10-D1* wild type (d1_wt; red open triangles) and transgenic (d1_mut; red filled triangles). Panel (a) Mesophyll cell wall thickness (T_CW_); (b) bundle sheath cell wall thickness (BS_CW_); (c) mesophyll surface area exposed to intercellular air spaces (S_mes_); (d) Chloroplast surface area exposed to intercellular air spaces (S_c_); (e) leaf thickness; (f) interveinal distance (IVD). Leaf anatomical traits were measured only for high-water treatment. *P*-values from pair-wise t-test are shown. Values indicate mean ± SE (*n*=5-6) along with replicate points (light gray dots).

### Impacts of modification of CW composition on leaf physiology and biochemistry

Both *Ubi_pro_-AT10 and OsAT10-D1* transgenics differed significantly from WT plants for several key leaf-level physiological traits as indicated by the significant genotype effects in a two-way ANOVA (Table 1–2 and S1). Specifically, *Ubi_pro_-AT10* transgenics exhibited a significantly greater g_m_ (120%, Fig. 3a) and A_net_ (22%, Fig. 3b) compared to WT (*P* < 0.001, Table 1). Transgenics also showed significantly greater g_sw_ (40%, Fig. 3c) compared to WT (*P* < 0.01, Table 1). The magnitude of increase in g_sw_ in transgenics was less compared to the 120% increase in g_m_, but greater compared to the 22% increase in A_net_. Consequently, though *Ubi_pro_-AT10* transgenics exhibited a significantly greater g_m_/g_s_ ratio (110%, *P* < 0.05, Table 1, Fig. 3d) compared to WT, WUE_i_ (that is, A_net_/g_sw_) remained unchanged (*P* > 0.1, Table 1, Fig. 3f). Also, we did not observe a significant genotype effect on C_i_/C_a_ ratio (*P* > 0.1, Table 1, Fig. 3e). In terms of biochemical traits, *Ubi_pro_-AT10* transgenics exhibited significantly greater J_max_ (20%, Fig. 4b) and A_max_ (14%, Fig. 4c) compared to WT (*P* < 0.1, Table 1). Whereas transgenics and WT exhibited similar V_cmax_ and leaf N content (*P* > 0.1, Table 1, Fig. 4a and d). There was a statistically significant genotype effect on stomatal, mesophyll and biochemical limitations (*P* < 0.01, Table 1) as*Ubi_pro_-AT10* transgenics exhibited lower relative stomatal (10%, Fig. 5a) and mesophyll limitations (60%, Fig. 5b), but greater biochemical limitations to A_net_ (50%, Fig. 5c) compared to WT. There were no significant genotype effects on leaf hydraulic traits, that is, K_leaf_, leaf water potential and K_x_ (*P* > 0.1, Table 1, Fig. 6a-c). However, K_ox_ tended to be lower (12%) in *Ubi_pro_-AT10* transgenics compared to WT (*P* = 0.09, Table 1, Fig. 6d).

**Figure 3.**
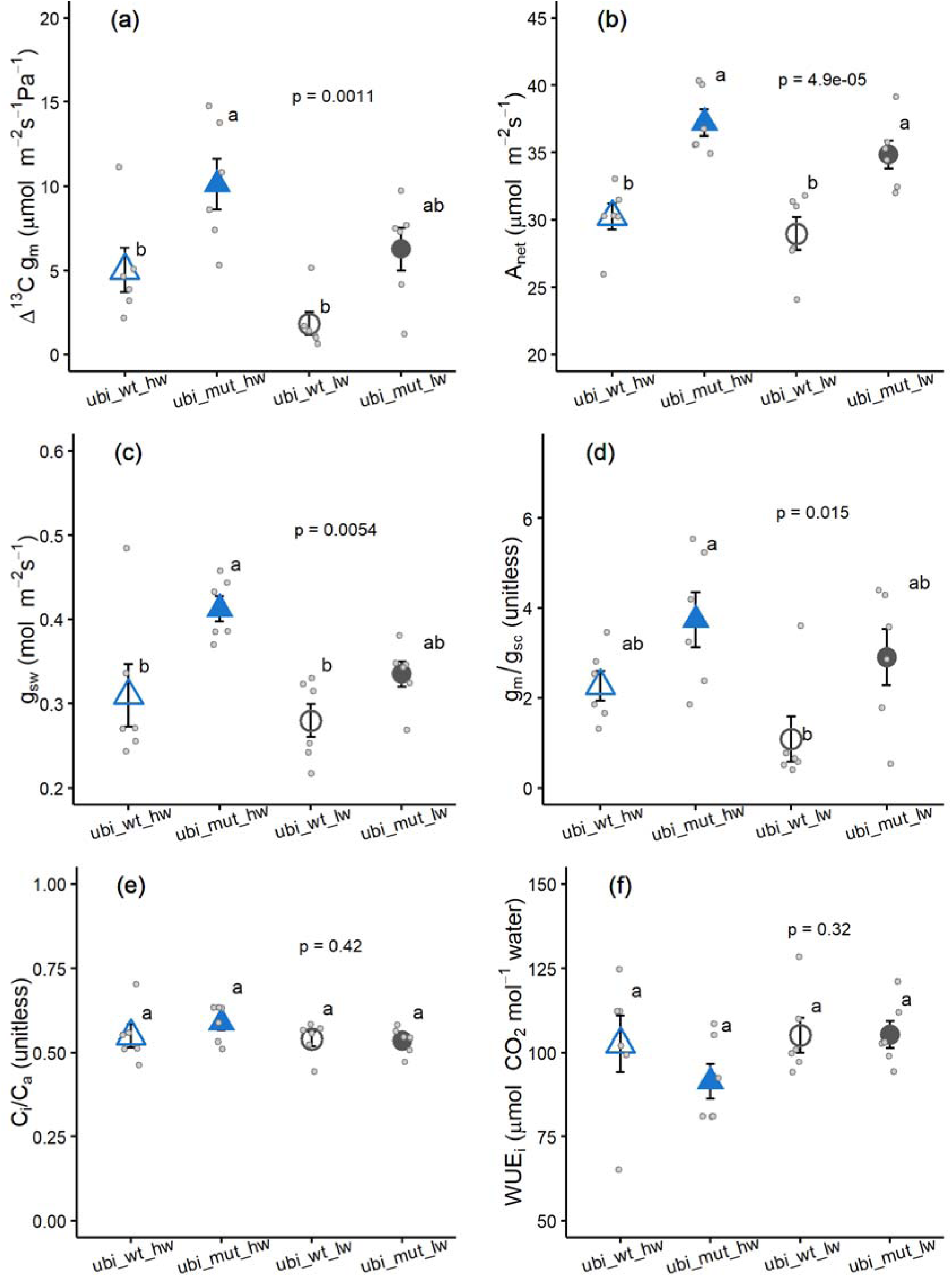
Leaf physiological traits for *Ubi_pro_-AT10* wild type (ubi_wt, open symbols) and transgenic (filled symbols) measured under high water (hw, blue triangles) and low water (lw, gray circles). (a) Mesophyll conductance to CO_2_ (Δ_13_C g_m_); (b) Net CO_2_ assimilation rates (A_net_); (c) Stomatal conductance to water (g_sw_); (d) ratio of g_m_ to g_sw_; (e) Ratio of leaf intercellular [CO_2_] to [CO_2_] in leaf chamber (C_i_/C_a_); and (f) Leaf-level water-use efficiency (TE_i_ = A_net_/g_sw_). *P*-values from one-way ANOVA and post-hoc Tukey’s test letters are shown. Values indicate mean ± SE (*n*=5-6) along with replicate points (light gray dots).

**Figure 4.**
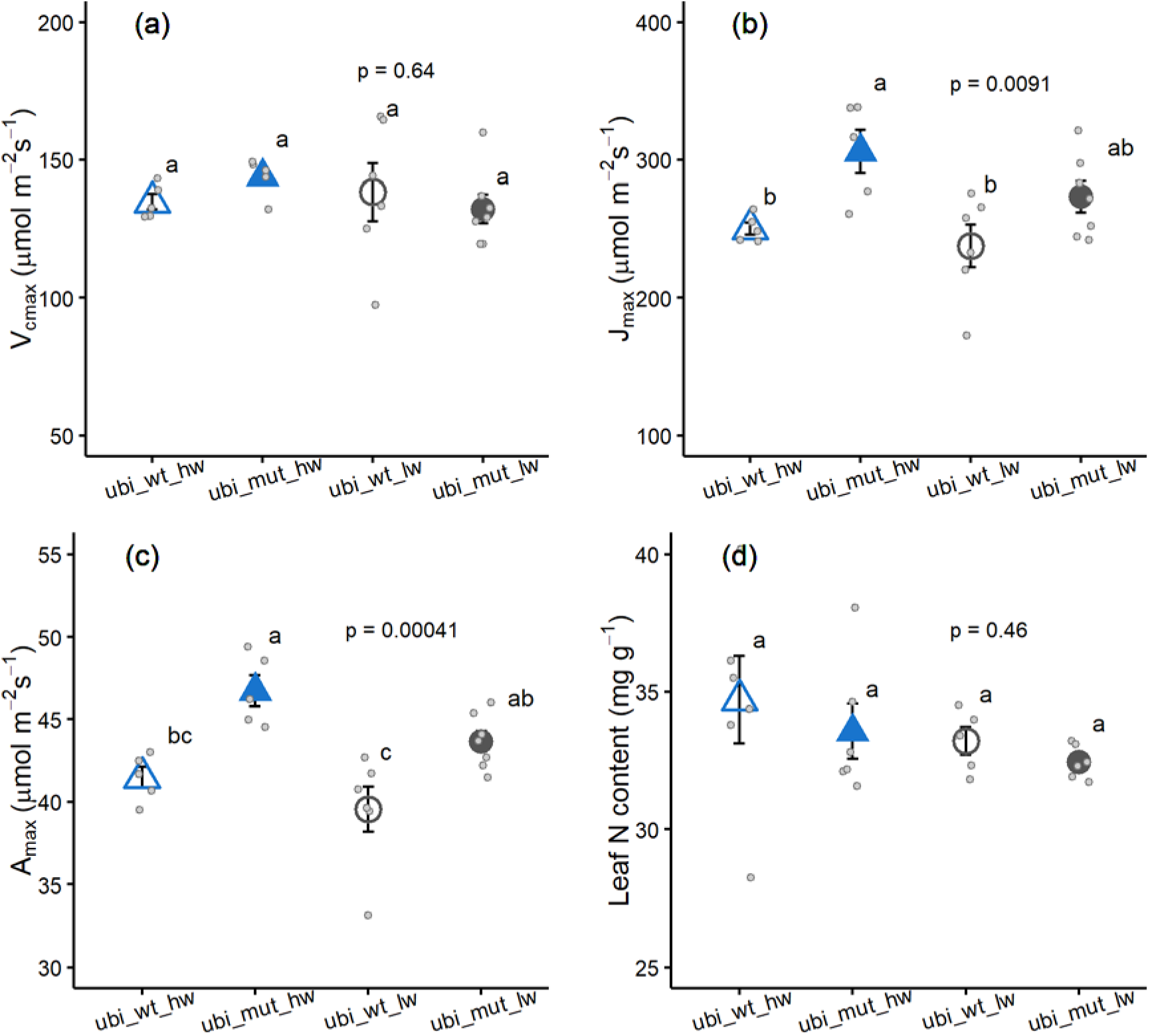
Biochemical traits for *Ubi_pro_-AT10* wild type (ubi_wt, open symbols) and transgenic (filled symbols) measured under high water (hw, blue triangles) and low water (lw, gray circles). (a) Maximum carboxylation capacity of Rubisco (V_cmax_); (b) Maximum electron transport rate (J_max_); (c) Maximum photosynthetic capacity at saturating *p*CO_2_ (A_max_) and (d) Leaf N content. *P*-values from one-way ANOVA and post-hoc Tukey’s test letters are shown. Values indicate mean ± SE (*n*=5-6) along with replicate points (light gray dots).

**Figure 5.**
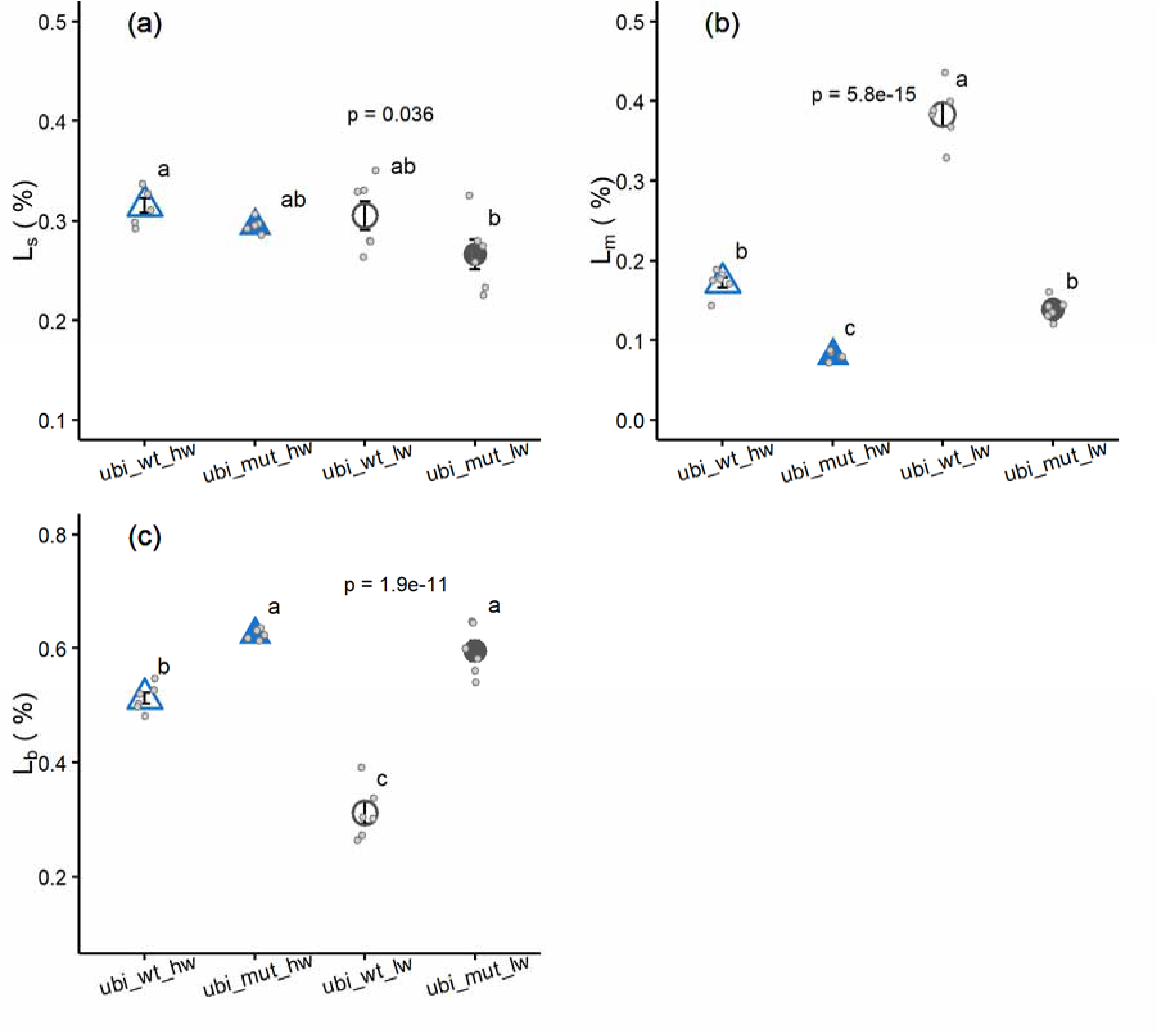
Relative limitations for *Ubi_pro_-AT10* wild type (ubi_wt, open symbols) and transgenic (filled symbols) measured under high water (hw, blue triangles) and low water (lw, gray circles). Panels show (a) stomatal (L_s_); (b) mesophyll (L_m_); and (c) biochemical (L_b_) limitations to photosynthesis. *P*-values from one-way ANOVA and post-hoc Tukey’s test letters are shown. Values indicate mean ± SE (*n*=5-6) along with replicate points (light gray dots).

**Figure 6.**
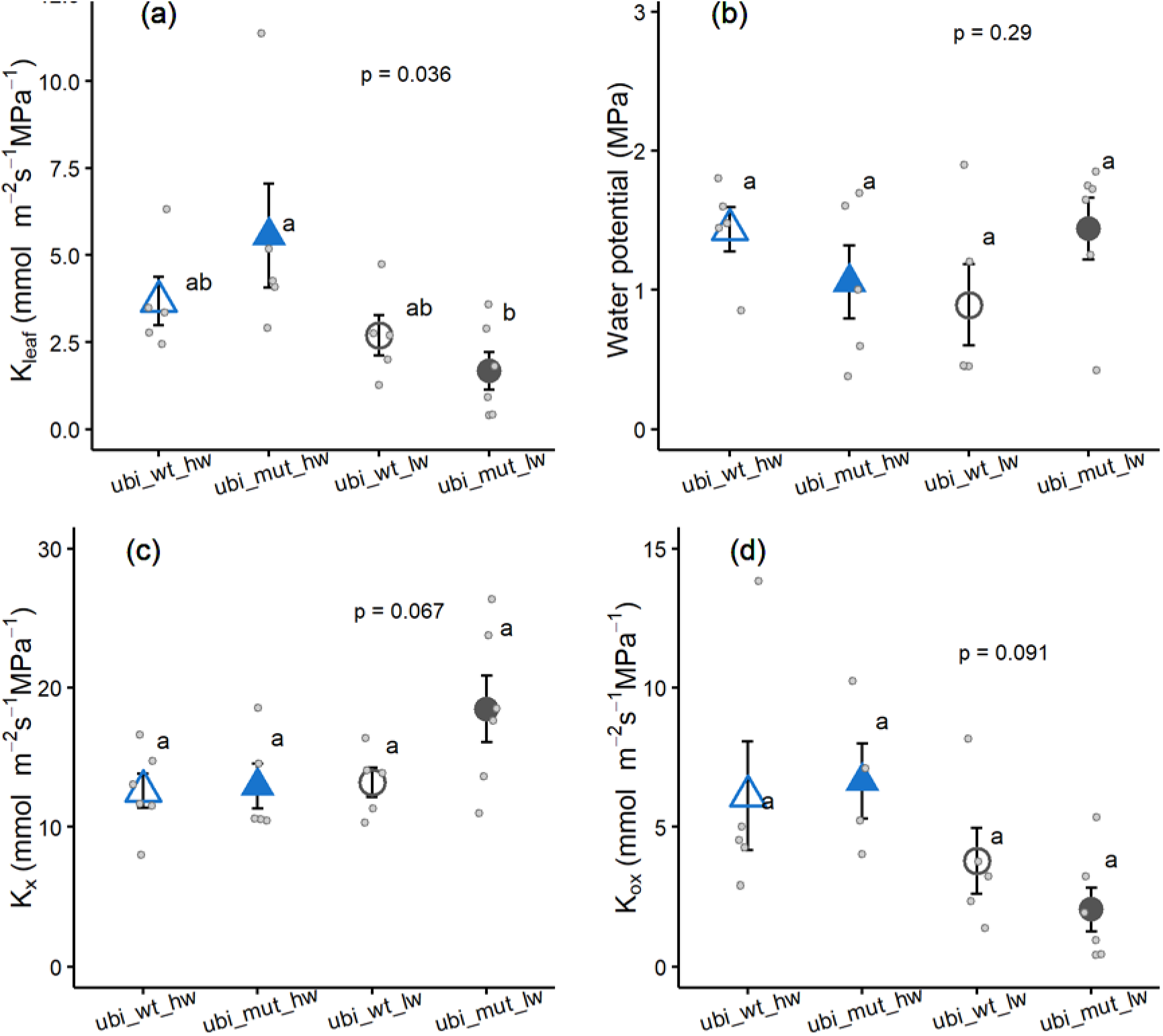
Leaf hydraulic traits for *Ubi_pro_-AT10* wild type (ubi_wt, open symbols) and transgenic (filled symbols) measured under high water (hw, blue triangles) and low water (lw, gray circles). (a) Leaf hydraulic conductance (K_leaf_); (b) Leaf water potential; (c) Xylem conductance (K_x_) and (d) outside xylem conductance (K_ox_). *P*-values from one-way ANOVA and post-hoc Tukey’s test letters are shown. Values indicate mean ± SE (*n*=5-6) along with replicate points (light gray dots).

**Table 2:** Summary of two-way ANOVA P-values and F-values for the plant-level traits measured on the rice *Ubi_pro_-AT10* wild type and transgenic lines at 50-60 days’ time point. *P*-values ≤ 0.1 are highlighted in bold. *P*-values ≤ 0.05 were considered as statistically significant and *P*-values ≤ 0.1 were considered as marginally significant.

| Abbreviations | Traits | Genotype |  | Water |  | Genotype x Water |  |
| --- | --- | --- | --- | --- | --- | --- | --- |
|  |  | *P*-value | *F*-value | *P*-value | *F*-value | *P*-value | *F*-value |
| AGB | Above-ground biomass | **0.005** | 9.88 | **< 0.001** | 115.07 | 0.624 | 0.25 |
| Leaf biomass | Biomass of transpiring surface | **0.039** | 4.93 | **< 0.001** | 161.93 | **0.048** | 4.47 |
| Stem biomass | Biomass of non-transpiring surfaces | **0.007** | 9.17 | **< 0.001** | 80.87 | 0.984 | 0.00 |
| Leaf width | Leaf width | **< 0.001** | 83.51 | **< 0.001** | 136.18 | 0.189 | 1.84 |
| LMA | Leaf mass per area | 0.373 | 0.83 | 0.745 | 0.11 | 0.170 | 2.05 |
| Leaf N content | Leaf nitrogen content | 0.325 | 1.02 | 0.218 | 1.63 | 0.854 | 0.04 |
| WUE<sub>canopy</sub> | Canopy-level water-use efficiency | **0.003** | 11.35 | 0.504 | 0.46 | 0.806 | 0.06 |
| Total water used | Total water-used during duration of experiment | 0.621 | 0.25 | **< 0.001** | 140.72 | 0.478 | 0.52 |

We did not observe a statistically significant genotype x water interaction effect on almost all the physiological and biochemical traits except for mesophyll and biochemical limitations to A_net_ (*P* < 0.01, Table 1). Particularly, the decrease in mesophyll limitations in *Ubi_pro_-AT10* transgenics compared to WT was more pronounced under lower water treatment (62%) compared to HW treatment (45%, Fig. 5b). Whereas the increase in biochemical limitations in *Ubi_pro_-AT10* transgenics compared to WT was more pronounced under LW treatment (100%) compared to HW treatment (30%, Fig. 5c).

Similar trends to those of *Ubi_pro_-AT10* lines were observed for key physiological traits in the *OsAT10-D1* lines. Particularly, we observed a significant genotype effect on key physiological traits wherein *OsAT10-D1* transgenics exhibited significantly greater g_m_ (65%), A_net_ (14%), g_sw_ (20%) and g_m_/g_sw_ (100%) ratio compared to WT (*P* < 0.05, Table S1, Fig. S2a-d). Whereas we did not observe a significant genotype effect on C_i_/C_a_ ratio and WUE_i_ (Fig. S2e and f). Also, we did not observe a significant genotype x water interaction effect on any of the physiological traits measured for *OsAT10-D1* lines (*P* > 0.1, Table S1).

### Impacts of modification of CW composition on plant-level traits

*Ubi_pro_-AT10* transgenics differed significantly from WT for key leaf and whole-plant traits as indicated by a statistically significant genotype effects (*P* < 0.05, Table 2, Fig. 7). In both water treatments, *Ubi_pro_-AT10* transgenics showed significantly greater AGB (12.5%, Fig. 7a), leaf biomass (7.74%, Fig. 7b), stem biomass (14.7%, Fig. 7c) and leaf width (14.3%, Fig. 7e) compared to WT. However, transgenics did not differ from WT in terms of LMA and leaf N content (Fig. 7d and f), nor in terms of total water used during the growth period (Fig. 7g). Consequently, there was a greater WUE_canopy_ (AGB/Total water used) in the transgenics compared to WT (8.8%, Fig. 7h). We did not observe significant genotype x water interaction effect for most leaf and whole-plant traits (*P* > 0.1, Table 2) except leaf biomass (*P* = 0.048, Table 2) which was significantly higher in *Ubi_pro_-AT10* transgenics than WT under lower water treatment (14.7%) but not under HW treatment.

**Figure 7.**
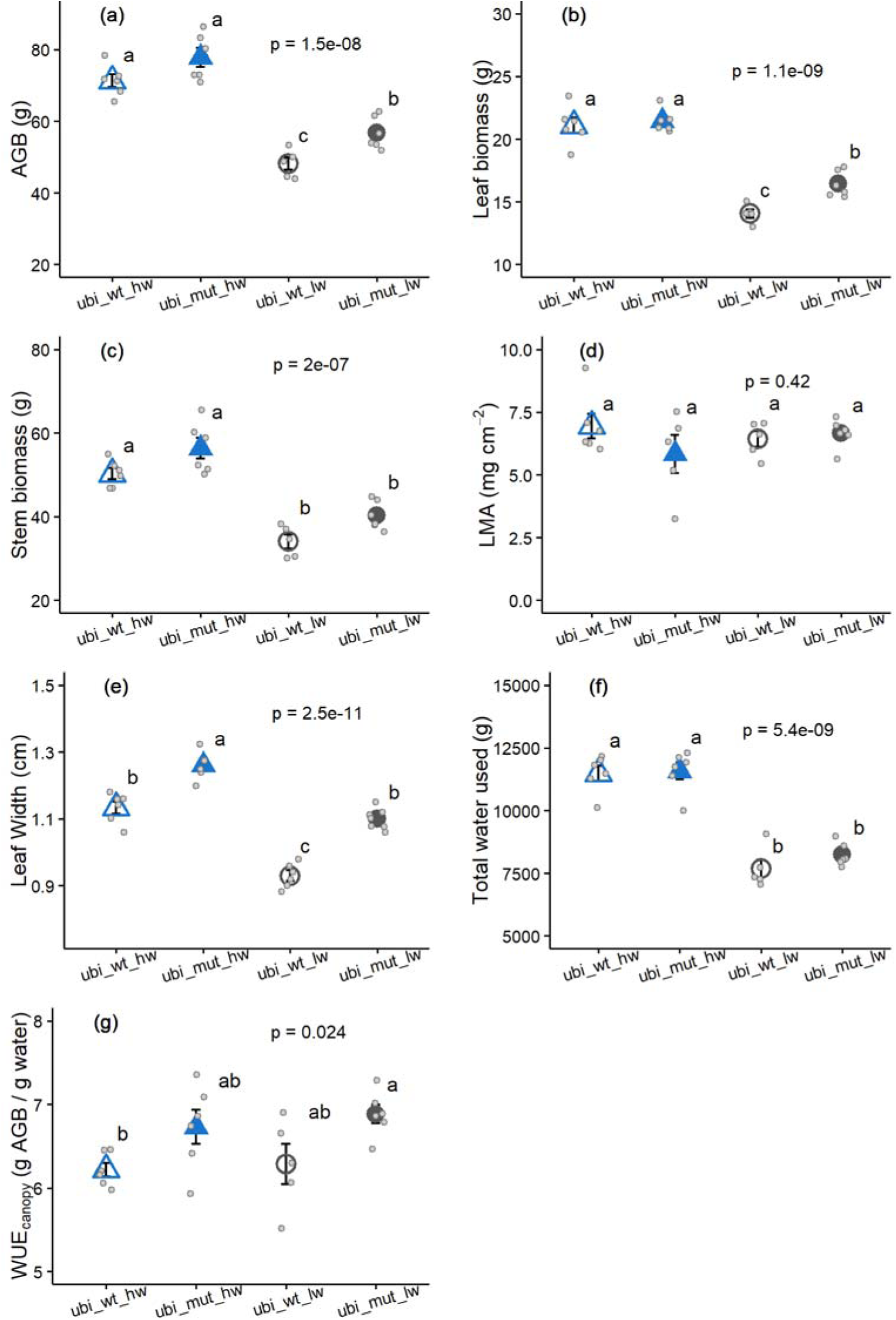
Leaf and plant-level traits for *Ubi_pro_-AT10* wild type (ubi_wt, open symbols) and transgenic (filled symbols) measured under high water (hw, blue triangles) and low water (lw, gray circles). (a) Total above-ground biomass (AGB); (b) total leaf or transpiring biomass; (c) total stem biomass or non-transpiring biomass; (d) Leaf mass per area (LMA); (e) Leaf width; (f) Total water used by plants during the experiment and (g) Canopy-level water-use efficiency (WUE_canopy_). *P*-values from one-way ANOVA and post-hoc Tukey’s test letters are shown. Values indicate mean ± SE (*n*=5-6) along with replicate points (light gray dots).

For *OsAT10-D1* lines we observed marginally significant genotype effects only on WUE_canopy_ (*P* = 0.073, Table S1, Fig. S3f), wherein *OsAT10-D1* transgenics exhibited greater WUE_canopy_ (4%) compared to WT. For other key traits like AGB, leaf biomass, stem biomass and total water used we observed a marginally significant genotype x water effect (*P* < 0.1, Table S1), wherein the *OsAT10-D1* transgenics showed greater AGB (10.5%, Fig. S3a), leaf biomass (7.5%, Fig. S3b) and stem biomass (12.4%, Fig. S3c), compared to WT only under LW treatment.

## Discussion

CW properties have been proposed as key targets for improving crop A_net_ and WUE_i_ (Lundgren and Fleming, 2020; Evans, 2021; Flexas et al., 2021; Pathare et al., 2022). However, little is known about how CW components affect g_m_, A_net_ and water-use in grasses—which comprise a majority of cereal crops. Here, we provide evidence that changes in CW hydroxycinnamic acid content (greater *p*-CA and lower FA) decrease CW thickness enhancing g_m_ and A_net_, without having a detrimental impact on leaf hydraulics and plant growth, even under LW stress.

### Changes in CW composition alters CW thickness and g_m_

Many recent studies report strong relationships of g_m_ with CW composition across diverse groups of trees and herbaceous dicot species, highlighting the importance of CWs for improving g_m_ and A_net_. However, there are only a few studies using transgenics to investigate the impacts of modification of CW composition on g_m_ and A_net_ in monocot grass species (Ellsworth et al., 2018). Specifically, Ellsworth *et al*., 2018 used rice plants with disruptions in CW mixed-linkage glucan (a polysaccharide unique to grasses) caused due to mutation in the cellulose synthase-like F-6 gene (CslF6) to assess the impacts of CW properties on g_m_ and other leaf physiological traits. The *cslF6* knockout plants exhibited significantly thinner CWs compared to WT but showed a substantial reduction in g_m_ (83%) and A_net_ (30-40%). This discrepancy between CW thickness and g_m_/*A*_net_ was attributed to lower CW effective porosity and increased effective path length. However, a significant portion of the reduction in g_m_ in the *cslF6* rice plants was also attributed to pleiotropic effects affecting other anatomical traits related to g_m,_ like S_c_, which was lower in the transgenics.

In the current study we explored how modification of CW hydroxycinnamic acids affect mesophyll CW thickness, g_m_ and other traits related to CO_2_-uptake and water-use. Hydroxycinnamates on arabinoxylan are another unique component of CWs of grasses and other recently evolved monocots, modification of which affects biomass properties (Bartley et al., 2013; Tian et al., 2021; Chandrakanth et al., 2023). Our data suggests that overexpression of *OsAT10*, which results in lower CW FA and greater *p*-CA content, leads to thinner CWs in the transgenics compared to WT plants. However, changes in FA and *p*-CA did not alter other anatomical traits known to influence g_m_ in rice like S_mes_, S_c_, leaf thickness and LMA (Fig.1) (Giuliani et al., 2013; Xiong and Flexas, 2018). Thus, in contrast to the results with *cslf6* disruption (Ellsworth et al., 2018), changes in g_m_ and A_net_ of the rice CW transgenics in the current study can be directly attributed to changes in CW composition and thickness. However, the coordinated changes in CW thickness and composition make it difficult to assess whether the increase in g_m_ was due to a decrease in CW thickness, changes in CW porosity, or a combination of both. Irrespective of the mechanism that led to the increase in g_m_, changes in CW hydroxycinnamic acid content in rice improve g_m_, i.e., the supply of CO_2_ to Rubisco.

### CW transgenics show greater g_m_ and A_net_, but not WUE_i_

Higher g_m_ in the rice CW transgenics was associated with a concurrent increase in A_net_ (Fig.3), thus supporting the previous reports that g_m_ is an important factor determining A_net_ in C_3_ species (Flexas et al., 2012; Barbour and Kaiser, 2016; Gago et al., 2020). However, the magnitude of the increase in A_net_ (22%) was low compared to increases in g_m_ (120%). This is because, besides diffusion of CO_2_, biochemical capacity is an important factor limiting A_net_ (Gago et al., 2020). Here we did not observe any significant differences in biochemical capacity (as indicated by leaf N content and V_cmax_) between the transgenics and WT. However, transgenics did show lower diffusional, but greater biochemical limitations compared to WT (Fig. 5). This suggests that the lesser magnitude of increase in A_net_ in the transgenics despite greater increases in g_m_ could be attributed to enhanced biochemical limitations in transgenics compared to WT. Alleviation of these biochemical limitations, for instance through changes in Rubisco properties and kinetics (Flexas et al., 2016; Carmo-Silva and Sharwood, 2023), could further increase A_net_ in the *OsAT10*-overexpression lines.

Though the rice transgenics exhibited greater g_m_ and A_net_ compared to wild type, we did not observe any concurrent increases in WUE_i_ (Fig. 3f). Because the CO_2_ diffusion pathway related to g_m_ is not the same as the pathway of water transpired out of the leaf through stomata, an increase in g_m_ is expected to increase CO_2_ concentrations at the site of fixation without a concurrent increase in g_sw_, thus increasing both A_net_ and WUE_i_ (Flexas et al., 2016). While there is some previous evidence supporting this (Sade et al., 2009; Flexas et al., 2013), studies on C_3_ species also report coordinated responses of g_m_ and g_sw_ such that WUE_i_ remains unchanged or even decreases (Giuliani et al., 2013; Barbour and Kaiser, 2016; Tomeo and Rosenthal, 2017). Such coordinated responses of g_m_ and g_sw_ were also observed in plants specifically engineered to alter g_m_ by changing CW composition or aquaporin expression (Hanba et al., 2004; Roig-Oliver et al., 2021). For instance, modification of CW composition in *Arabidopsis* led to simultaneous decreases in g_m_, g_sw_ and A_net,_ resulting in no differences in WUE_i_ between transgenics and WT plants (Roig-Oliver et al., 2021). We observed the same pattern in the current study, with increases in g_m_ in the transgenics also being associated with a concomitant increase in g_sw_. Together these results suggest that manipulation of g_m_ can increase A_net_, but concomitant changes in g_sw_ can make a simultaneous increase in A_net_ and WUE_i_ difficult. Thus, to achieve the goal of greater A_net_ and WUE_i_, biotechnological manipulations must induce increases in g_m_, whilst maintaining or even decreasing g_sw_ (Leakey et al., 2019). Furthermore, our study supports previous reports (Roig-Oliver et al., 2021) that changes in CW properties can affect g_sw_. Whether such relationships observed between CWs and g_sw_ result from direct effects of CW properties on stomata (Jones et al., 2003) or an indirect effect of co-adjustment between g_m_ and g_sw_ (Flexas et al., 2012) needs to be investigated.

### Changes in CW properties do not alter K_leaf_

In the current study, we also investigated how modification of CW properties and thickness affect the water transport inside leaves (K_leaf_) and the coordination of K_leaf_ with g_m_, g_sw_ and A_net_. Just like g_m_, K_leaf_ is extremely complex and influenced by several leaf anatomical features (Buckley et al., 2015). However, unlike g_m_, the impacts of different leaf anatomical features on K_leaf_ and primarily K_ox_, are less studied. The rice CW transgenics in the current study showed significantly lower CW thickness and altered CW composition without concurrent changes in other anatomical traits known to influence K_leaf_ like IVD, thickness, bundle sheath CW thickness, S_mes_ and stomatal density (Fig. 1). This provided us with the unique opportunity to test the impacts of modification of mesophyll CWs on K_leaf_ and K_ox_. Despite a significant decrease in mesophyll CW thickness, we observed no changes in K_leaf_ and K_ox_ in the transgenics. The apoplastic pathway via mesophyll CWs is often assumed to be the major pathway of water movement outside the xylem, based on which it has been hypothesized that thinner CWs would lead to lower resistance to water movement by decreasing the outside xylem apoplast pathlength, thus leading to lower K_leaf_ and K_ox_ (Buckley et al., 2015). Our results contrast these expectations and suggest that mesophyll CW thickness may not be the only factor determining K_leaf_ and K_ox_ in rice. Instead, symplastic water movement could have major influences on K_ox_, as suggested previously (Sade et al., 2014; Buckley et al., 2015).

Similar to the coupling between g_m_ and g_sw_, previous studies also demonstrate a coupling of g_m_ and g_sw_ with K_leaf_. For example, in diverse C_3_ species and under different environmental conditions, g_sw_ and K_leaf_ were correlated, presumably because of the common pathway of movement shared by water and CO_2_ via stomata and cuticle (Boyer, 2015; Scoffoni et al., 2016) and via the mesophyll tissue inside leaves (Flexas et al., 2013; Xiong et al., 2017). Such coordinated changes in g_m_, g_sw_ and K_leaf_ have also been demonstrated in studies that use transgenics to modify these conductances (Sade et al., 2014). However, in the current study, we did not observe changes in K_leaf_ and K_ox_ despite changes in g_sw_, g_m_ and A_net_. Such decoupling of leaf hydraulic and photosynthetic traits has been reported previously for some environmental conditions (Li et al., 2015) and plant species (Pathare et al., 2020) and implies that improving crop g_m_, A_net_ and WUE_i_, whilst maintaining K_leaf_, is possible.

### Changes in CW properties affect AGB and WUE_canopy_

CWs play a major role in maintaining structural integrity and drought tolerance capacity of plants (Keegstra, 2010). Consequently, one could expect that any CW-related improvement in g_m_ and A_net_ will have detrimental effects on plant tolerance to water-stress and/or any other fundamental functions performed by CWs in plants. Here, in addition to the greater g_m_ and A_net_, rice CW transgenics also exhibited a small (12.5%) but statistically significant increase in AGB under both HW and LW treatments (Fig. 7). The increase in AGB was not associated with a concurrent increase in canopy water use, because of which we also observed a significant increase in WUE_canopy_ in CW transgenics. Our results thus demonstrate that modification of CW hydroxycinnamic acid content improves g_m_ and A_net_ without having a detrimental effect on plant growth, even under LW stress.

## Conclusions

In summary, our results demonstrate that alteration of CW hydroxycinnamic acid content (greater *p*-CA and lower FA) decreased mesophyll CW thickness, and enhanced g_m_ and A_net_ in rice. However, a concomitant increase in g_sw_ canceled out the expected benefit of increased A_net,_ resulting in a lack of changes in WUE_i_. Despite no changes in WUE_i_, rice transgenics exhibited greater AGB and WUE_canopy_, with a greater increase under LW availability. The alteration of rice CW hydroxycinnamic acid content and mesophyll CW thickness did not influence leaf hydraulic traits. Overall, our results indicate that modification of CW hydroxycinnamic acid content, a unique and important feature of grass CWs, can influence mesophyll CW thickness and leaf and plant-level CO_2_-uptake and water-use without having detrimental impacts on plant growth even under LW stress. Integrating such increases in g_m_ and A_net_, through changes in CW hydroxycinnamic acid content, with efforts to increase photosynthetic efficiency (Carmo-Silva and Sharwood, 2023) and decrease g_sw_ (Leakey et al., 2019) could help achieve substantial increases in photosynthesis, biomass, and WUE under both well-watered and water limited conditions in monocot C_3_ crops.

## Acknowledgements

This work was supported by the Division of Chemical Sciences, Geosciences, and Biosciences, Office of Biological and Environmental Research in the DOE Office of Science (DE-SC0018277 and DE-SC0023160), the National Science Foundation (Major Research Instrumentation grant no. 0923562), and USDA-NIFA (Hatch project #1015621). We are also grateful to the Core Facility Center “Cell and Molecular Technologies in Plant Science” of Komarov Botanical Institute (St.-Petersburg, Russia) and Franceschi Microscopy and Imaging Center at Washington State University (Pullman, USA) for the use of its facilities and staff assistance. We would also like to thank Charles A. Cody for help in plant growth management. The authors declare that they have no conflicts of interest.

## Author contributions

V.S.P., L.E.B., and A.B.C. designed the experiment. V.S.P., R. P., B.V.S., A. J. A., and N.K. performed the measurements and analyzed the data. V.S.P. led the writing with constructive inputs from all authors.

## Data availability

Data available on request from the authors.

